# Pan-cancer estimation of prognostic immune biomarkers from histopathology using artificial intelligence

**DOI:** 10.64898/2026.10.06.757083

**Authors:** Yijun Li, Abdul Rahman Diab, Abdallah Lamane, James R. Lindsay, Scott J. Rodig, Rafael A. Irizarry, William Lotter

## Abstract

Multiplex immunofluorescence (mIF) assays such as ImmunoProfile provide standardized quantification of prognostic immune markers but are costly and not routinely used in clinical settings. We developed a weakly-supervised AI model, HistoProfile, to estimate intratumoral cell densities of key immune biomarkers (CD8, FOXP3, PD-1, and PD-L1) and the PD-L1 tumor proportion score from routine H&E-stained whole-slide images. Trained on a large pan-cancer cohort with paired H&E and mIF data, HistoProfile discriminated biomarker-high from biomarker-low tumors in held-out samples and tumor-specific subgroups (AUROCs of 0.78-0.89 across biomarkers). Critically, the estimated biomarkers preserved prognostic stratification in downstream survival analyses, and showed cancer-specific survival associations across TCGA cohorts where mIF is unavailable. These findings support scalable H&E-based estimation of key immune biomarker measurements for large-scale immune profiling and risk stratification.

## Introduction

The tumor immune microenvironment (TIME) critically influences clinical outcomes and treatment response across cancer types^1,2^. Cytotoxic CD8^+^T cells, regulatory T cells (FOXP3^+^), and immune checkpoints such as PD-1 and PD-L1 are routinely studied as prognostic and predictive biomarkers, and composite immune scores have been proposed for patient risk stratification^3–6^. Emerging imaging techniques such as multiplex immunofluorescence (mIF) enable simultaneous quantification of these biomarkers within the TIME^7–11^. While commonly performed in research settings, a recent mIF study in a clinical cohort of 2,023 patients demonstrated that intratumoral densities of several immune biomarkers (CD8, FOXP3, PD-1, PD-L1) were associated with patient survival across 14 major cancer types^5^. However, the widespread implementation of this assay, known as ImmunoProfile, and mIF more generally is limited by its cost and expertise needed to execute expert-in-the-loop mIF workflows.

By contrast, hematoxylin and eosin (H&E) staining is widely performed as part of standard pathology workflows. Recent studies have explored the potential of predicting mIF-derived features directly from digital H&E images using artificial intelligence (AI)^12–14^. Valanarasu et al. developed GigaTIME, an image-to-image translation approach to generate virtual mIF images from paired H&E regions of interest (ROIs)^12^. GigaTIME used a UNet-based^15^ architecture trained on co-registered images from 21 H&E-stained lung adenocarcinoma slides. Channel-wise average pixel intensities from the resulting virtual 21-channel mIF images were used as inferred features to identify numerous biomarker associations across pan-cancer H&E whole-slide image (WSI) cohorts. These findings demonstrate the potential to recover mIF-related information from routine histopathology, but important gaps remain. In particular, approaches are needed that directly estimate biologically interpretable, quantitative features, such as intratumoral cell densities, using larger and more diverse cohorts, ideally without requiring technically challenging mIF/H&E co-registration. Furthermore, it remains unknown whether H&E-derived estimates can recover not only the quantitative values of mIF-measured biomarkers but also their downstream prognostic characteristics.

Here, we develop HistoProfile, an AI model to estimate mIF-derived immune biomarkers directly from routine H&E WSIs (**Fig. 1**). Developed using a pan-cancer cohort of 2,000 paired but unregistered mIF and H&E WSIs, the model uses a multiple instance learning framework to estimate the quantitative measurements from the original ImmunoProfile workflow using H&E alone. We evaluate whether the H&E-derived immune profiles generalize across tumor types, preserve clinically meaningful risk stratification, and provide spatially interpretable signals that align with, and extend beyond, predefined mIF sampling regions. We further assess generalizability in an independent external mIF dataset and prognostic associations in 11 TCGA cohorts, supporting the potential of H&E-based immune profiling to broaden access to a CLIA-validated assay and facilitate large-scale retrospective studies of key immune biomarkers.

**Figure 1.**
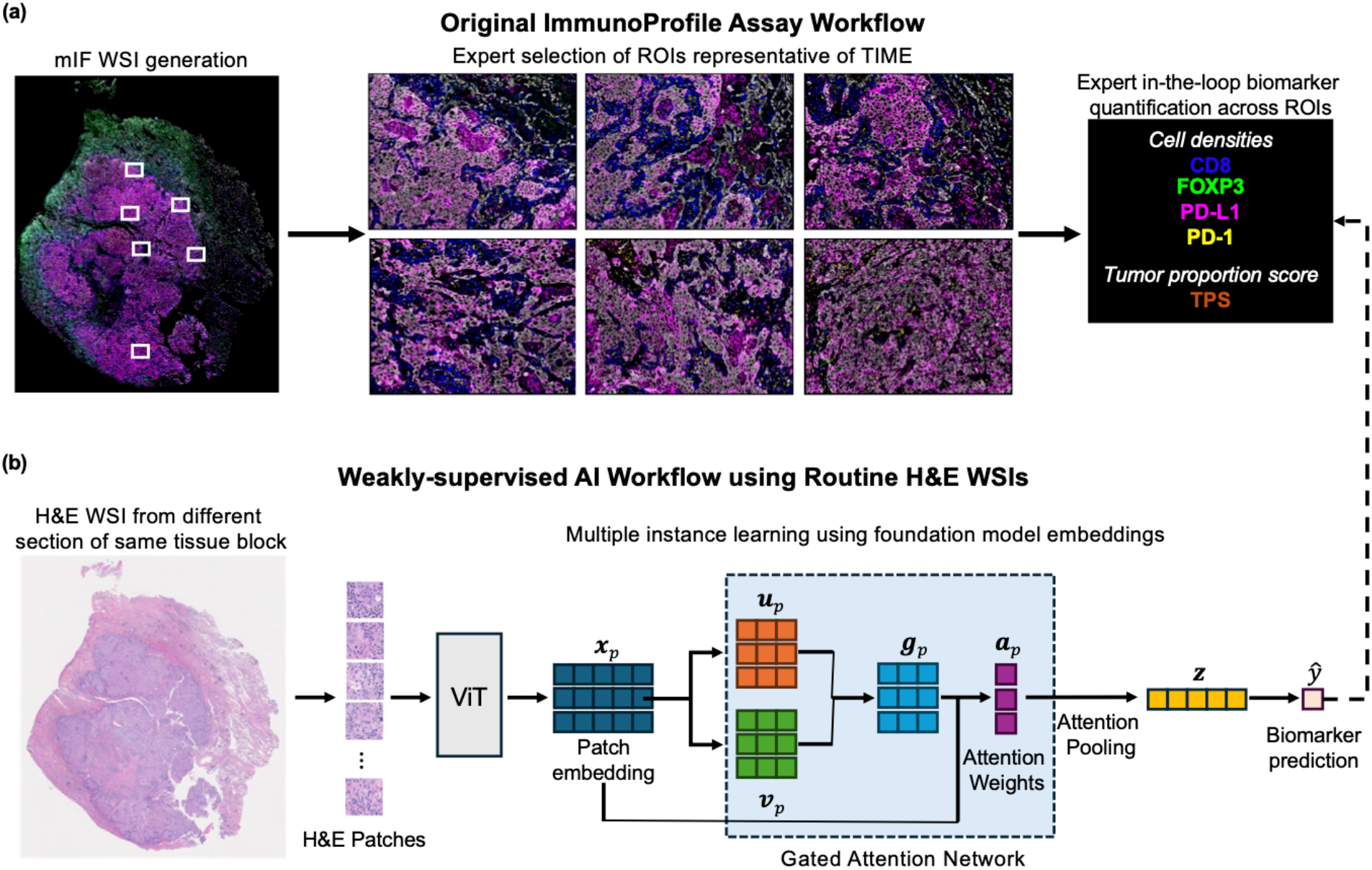
Overview of the HistoProfile framework for estimating immune biomarkers from routine H&E whole-slide images. (a) Original ImmunoProfile assay workflow. Multiplex immunofluorescence (mIF) whole-slide images (WSIs) are generated, followed by expert selection of regions of interest (ROIs) representative of the tumor immune microenvironment (TIME). Expert-in-the-loop analysis across ROIs quantifies the densities of CD8, FOXP3, PD-L1, and PD-1 positive cells and the PD-L1 tumor proportion score (TPS). (b) Weakly supervised AI workflow. HistoProfile follows a multiple instance learning workflow wherein an H&E-stained WSI from a different section of the same tissue block is divided into patches, which are encoded by a foundation model vision transformer (ViT) to obtain patch-level embeddings *x*_*p*_. A gated-attention network transforms the embeddings and assigns attention weights *a*_*p*_; attention pooling then aggregates the weighted patch representations into a slide-level representation *z*. This representation is used to estimate the corresponding slide-level biomarker value *ŷ*, with the measurements obtained from the ImmunoProfile assay serving as training targets. A separate multiple instance learning network is trained for each biomarker.

## Results

### ImmunoProfile Cohort

Model development used the Dana-Farber Cancer Institute ImmunoProfile dataset, a prospective pan-cancer cohort collected in routine clinical practice through a CLIA-validated mIF workflow^5,16^. This cohort includes 14 major cancer types and spans both primary and metastatic specimens as well as low- and high-stage disease, consisting of a real-world, unselected oncology population. For each case, H&E and mIF sections were generated from the same FFPE tissue block. The mIF assay stains for four immune biomarkers (CD8, FOXP3, PD-1, PD-L1), a tumor marker, and DAPI. Following a standardized workflow, intratumoral cell densities for each immune biomarker and the PD-L1 tumor proportion score (TPS) were quantified over expert-selected regions of interest from the mIF WSI. After applying exclusion criteria related to insufficient tissue and data missingness (see **Methods**), the final cohort consisted of 2000 cases from 2000 patients (**Supplementary Table 1**). The cohort was split into training (60%), validation (10%), and test (30%) sets while stratifying for major cancer types and cancer stage.

### HistoProfile: H&E-based multiple instance learning model for estimating mIF-derived immune biomarkers

HistoProfile uses a multiple instance learning framework to estimate ImmunoProfile biomarker measurements from H&E WSIs (**Fig. 1**). Each WSI is divided into image patches, patch embeddings are computed using a foundation model, and attention-based multiple instance learning (ABMIL)^17^ learns which regions are most informative for estimating the mIF-derived biomarker measurements (**Fig. 1**). A separate ABMIL head is fit for each metric (CD8, FOXP3, PD-1, PD-L1 cell densities and PD-L1 TPS), providing a weakly supervised framework that directly estimates biologically and clinically meaningful measurements without relying on H&E–mIF alignment. Moreover, whereas the ImmunoProfile assay quantifies biomarkers from selected ROIs to improve throughput, HistoProfile generates estimations using the entire H&E WSI, potentially enabling a more global and less operator-dependent TIME characterization. For each metric, the model was trained on percentile-normalized values across the ImmunoProfile train split using mean absolute error.

### HistoProfile accurately estimates biomarker measurements across cancer types

We first evaluated how well HistoProfile could discriminate between patients with high versus low values for each biomarker measurement, mimicking previous clinical studies of patient stratification^5,12^. In the pan-cancer test set, receiver operating characteristic (ROC) curves for estimated CD8, FOXP3, PD-L1, PD-1 and TPS demonstrated high area under the curve (AUC) values for all metrics, with a range of 0.78-0.89 AUROC (**Fig. 2a**). Subgroup analyses restricted to individual tumor types—breast cancer, colorectal cancer, non-small-cell lung cancer (NSCLC) and other cancers—showed that this discriminative performance was maintained across cancers, with consistently high AUCs for each biomarker within each subgroup.

**Figure 2.**
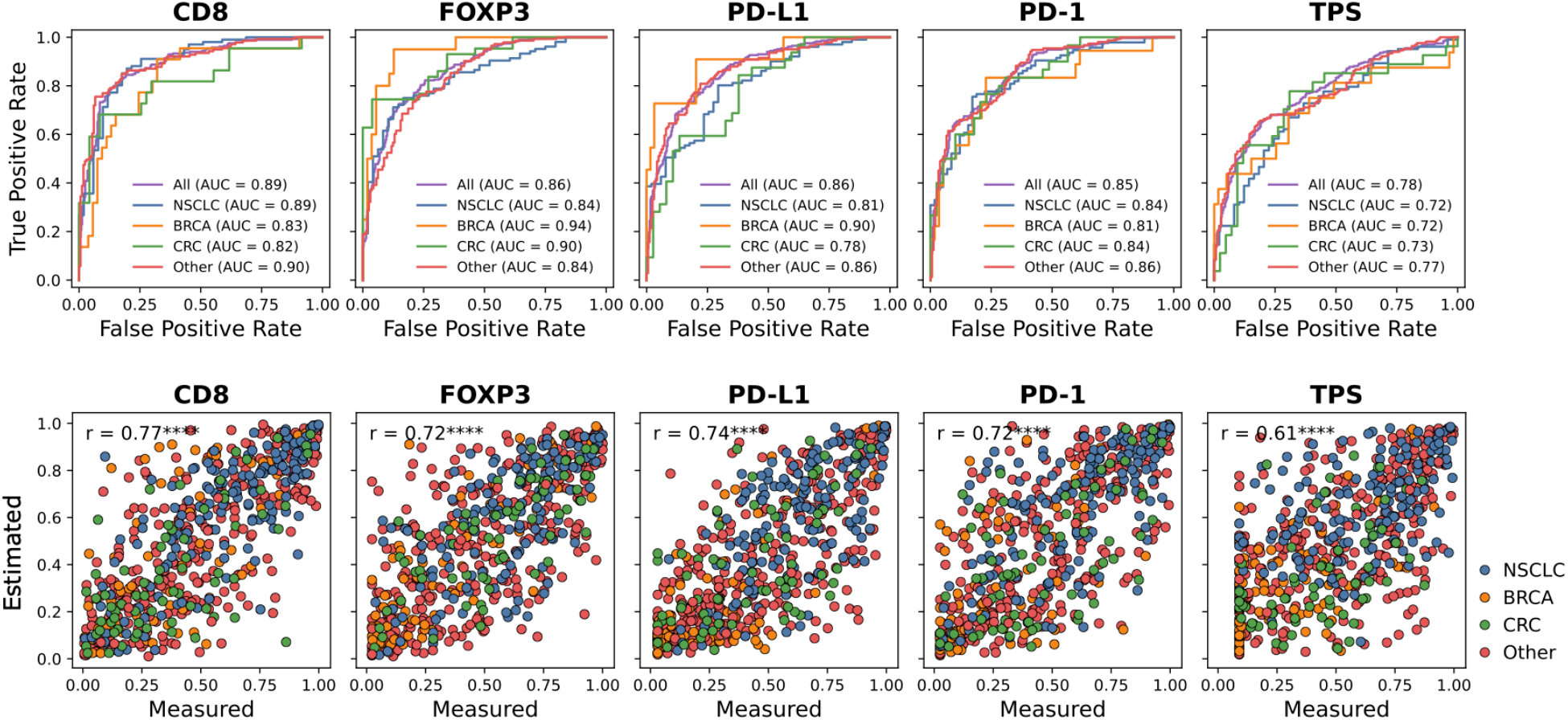
HistoProfile accurately estimates immune biomarkers in the ImmunoProfile pan-cancer cohort. (a) Receiver operating characteristic (ROC) curves evaluating HistoProfile estimations of high versus low CD8, FOXP3, PD-L1, PD-1, and TPS in the ImmunoProfile test split, shown for the entire pan-cancer split (“All”) and separately for non-small cell lung cancer (NSCLC), breast cancer (BRCA), colorectal cancer (CRC), and other cancer types. (b) Correspondence between measured and estimated percentile-normalized biomarker values. Each point represents one sample and is colored by cancer type (**** *P*<0.0001).

As HistoProfile was trained to estimate continuous, percentile-normalized metrics, we also examined the agreement between estimated and measured continuous values. Scatter plots of true versus estimated values showed strong and statistically significant Pearson correlation for each biomarker with a range of 0.61-0.77 (**Fig. 2b**). In addition, HistoProfile estimates recapitulated both pairwise correlations between biomarkers and pairwise differences in their measured levels, supporting the model’s ability to distinguish among individual biomarkers rather than primarily capturing a general pattern of immune infiltration (**Supplementary Fig. 1**).

Together, these results indicate that HistoProfile can approximate the CLIA-validated ImmunoProfile biomarker measurements from H&E WSIs across diverse cancer types.

### HistoProfile-estimated biomarkers retain prognostic information

We next asked whether the HistoProfile estimates retain the prognostic signal previously observed for ImmunoProfile measurements. Using Cox proportional hazards models adjusted for cancer stage, we compared the concordance indices (C-indices) for overall survival achieved when including either measured or estimated biomarker values, both in the overall pan-cancer test set and within individual cancer types. Across CD8, FOXP3, PD-L1, PD-1 and TPS, the HistoProfile estimates achieved C-indices that were consistently similar to those using the mIF-derived measurements (**Fig. 3a**). In the pan-cancer test split, CD8 density was the biomarker most associated with overall survival for both measured and estimated values (0.628, 95% CI [0.579, 0.674] vs. 0.624, 95% CI [0.565, 0.661], respectively), with TPS showing the least association (0.536, 95% CI [0.460, 0.563] vs. 0.511, 95% CI [0.481, 0.583], respectively). Kaplan-Meier curves stratified by estimated biomarker values also showed separations comparable to those based on measurements (**Fig. 3b**). Together, these findings suggest that the HistoProfile-estimated biomarker values preserve the prognostic information present in the measured biomarkers in the ImmunoProfile cohort.

**Figure 3.**
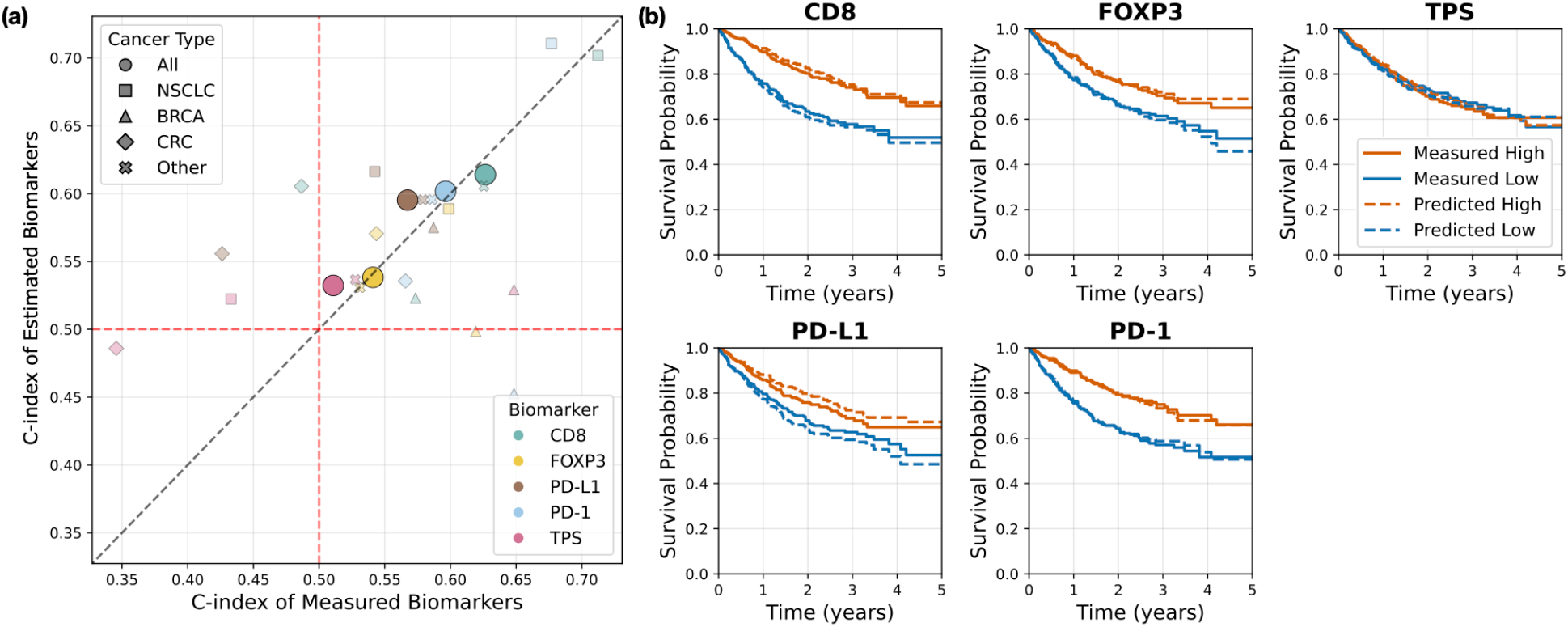
HistoProfile-estimated biomarkers retain prognostic characteristics of the measured biomarkers in the ImmunoProfile cohort. (a) Comparison of the C-index for overall survival of estimated versus measured biomarkers in the ImmunoProfile test split. (b) Kaplan-Meier curves of each biomarker stratified by median split separately based on measured and estimated values.

### Attention maps recapitulate mIF ROIs and highlight additional regions

To investigate the histologic features contributing to HistoProfile estimates, we visualized model attention across representative NSCLC samples from the ImmunoProfile test split with high and low measured CD8 densities (**Fig. 4**). In the CD8-high sample (**Fig. 4a**), regions receiving high model attention were broadly distributed across the tumor region and overlapped spatially with the regions annotated on the mIF image for biomarker quantification. H&E patches with the highest attention frequently contained prominent lymphocyte infiltration, consistent with the elevated measured CD8 density, and HistoProfile’s normalized CD8 density estimate was similar to the measured value. Conversely, the CD8-low sample exhibited more limited and heterogeneous regions of high attention (**Fig. 4b).**

**Figure 4.**
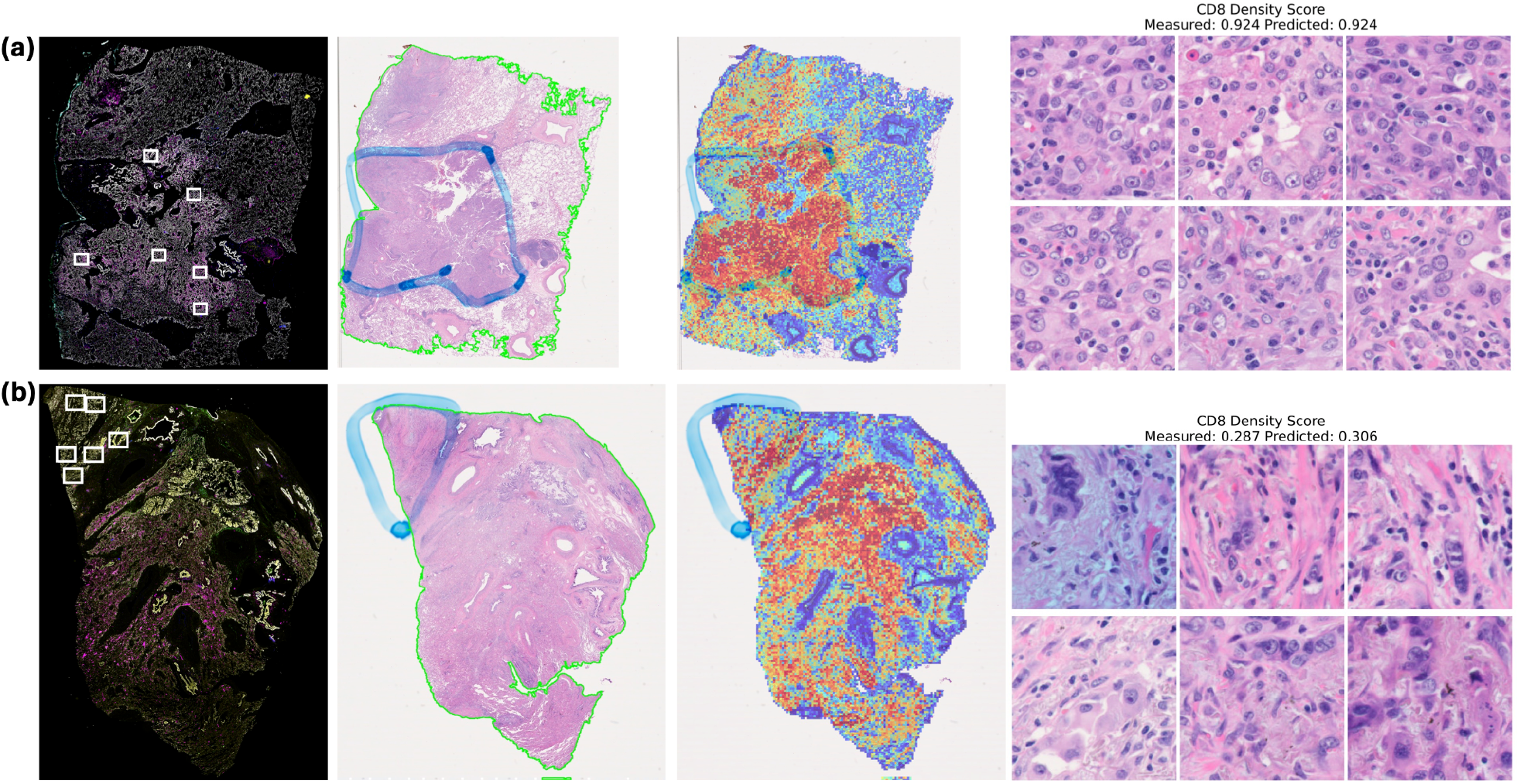
Visualization of HistoProfile attention patterns for representative samples with high and low CD8 density. **(a, b)** Representative samples with high (a) and low (b) normalized CD8 density scores. From left to right, each row shows the multiplex immunofluorescence (mIF) whole-slide image with expert-selected regions of interest indicated by white boxes, the corresponding H&E whole-slide image with the analyzed tissue region outlined, the HistoProfile attention map, and the most highly attended H&E patches. Warmer colors in the attention map indicate higher model attention. Measured and HistoProfile-estimated CD8 density scores are reported above the patch panels. Both examples are from NSCLC primary tumor specimens.

High attention patches showed comparatively sparse lymphocyte infiltration, and the estimated normalized CD8 density remained close to the measured value. Examples for the remaining biomarkers demonstrate consistent findings (**Supplementary Figs. 3-6)**. Furthermore, the attention maps also highlighted additional areas outside the originally sampled ROIs, coinciding with high-PanCK tumor regions. Together, these visualizations suggest that HistoProfile prioritizes biologically relevant tissue regions when generating slide-level estimations.

### HistoProfile preserves prognostic biomarker information in an independent colorectal cancer cohort

We next evaluated whether the survival-associated information captured by HistoProfile generalized to an independent colorectal cancer cohort acquired using a different imaging platform and processing pipeline. The ORION-CRC dataset provides matched mIF and H&E images, together with progression-free survival data for 40 patients^18^. We applied HistoProfile to the H&E WSIs to estimate CD8, FOXP3, PD-L1, and PD-1 intratumoral cell densities and PD-L1 TPS and compared their prognostic performance with the corresponding values measured by mIF (**Fig. 5**).

**Figure 5.**
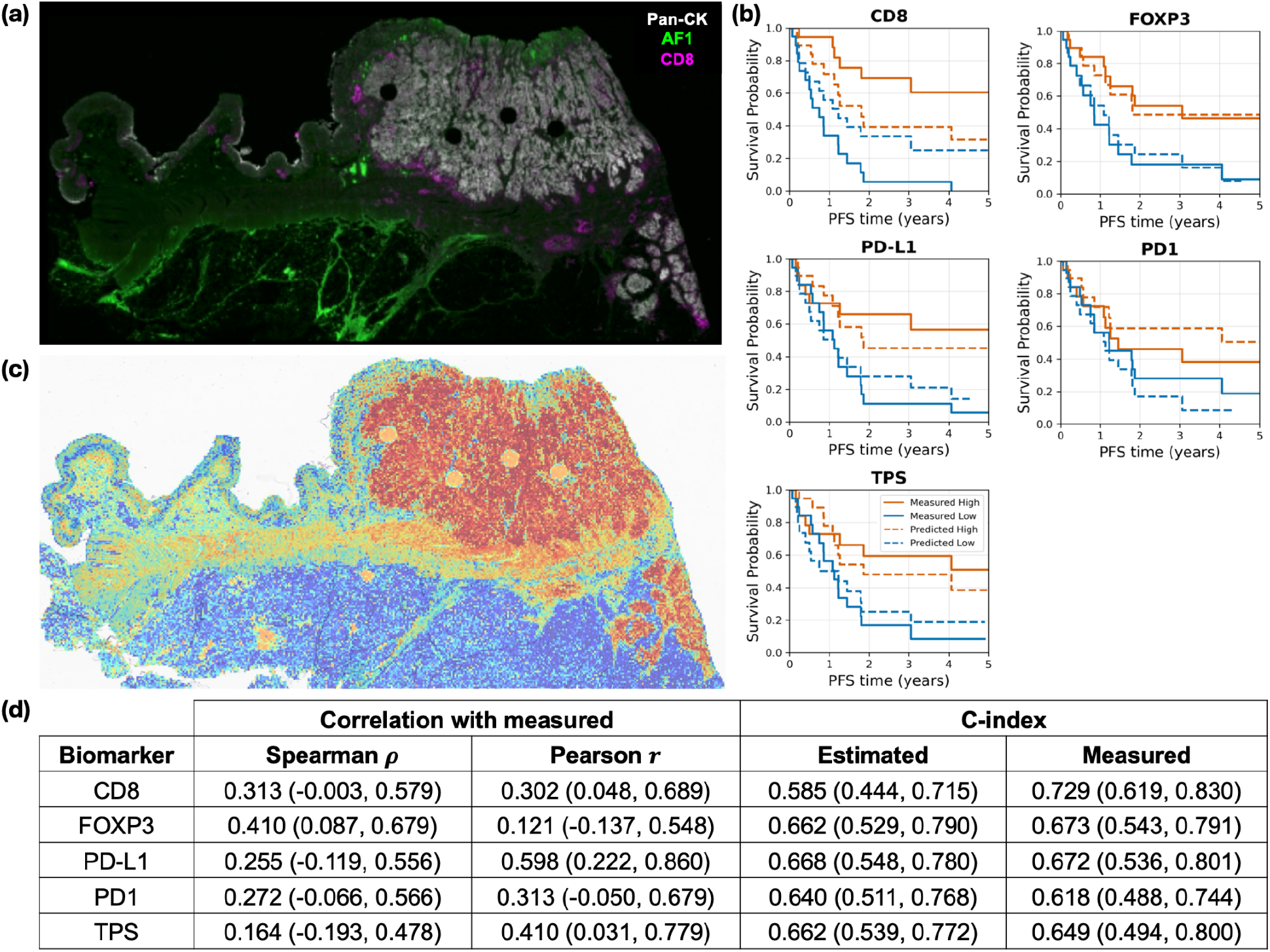
External validation of HistoProfile in the independent ORION-CRC cohort. **(a)**Representative multiplex immunofluorescence (mIF) whole-slide image showing pan-cytokeratin (Pan-CK), autofluorescence channel 1 (AF1), and CD8. **(b)** Kaplan–Meier survival curves comparing patient stratification based on measured and HistoProfile-estimated CD8, FOXP3, PD-L1, PD-1, and TPS values. High and low groups were defined using the respective cohort medians; solid lines represent groups defined using measured values, and dashed lines represent groups defined using estimated values. **(c)** HistoProfile attention map for the representative tissue section shown in **(a)**, with warmer colors indicating higher model attention. **(d)** Correlation between measured and HistoProfile-estimated biomarkers and concordance indices (C-indices) for progression-free survival.

Across all five studied biomarkers, stratification using HistoProfile estimates reproduced the direction of the survival separation obtained using measured values (**Fig. 5b**). C-index comparisons further supported this finding (**Fig. 5d**), where HistoProfile-estimated biomarkers achieved similar C-indices as mIF-derived measurements. Correlations between estimated and measured biomarkers were positive but lower in magnitude than in the ImmunoProfile cohort, which may be partly attributable to differences in biomarker quantification (see Methods). A CD8-high example is displayed in **Fig. 5a**, wherein HistoProfile’s attention map aligns with regions of high CD8 expression. Collectively, these results demonstrate that HistoProfile can generalize to an external cohort, preserving much of the prognostic discrimination present in the measured biomarkers.

### HistoProfile biomarker estimates show cancer-specific survival associations in external TCGA cohorts

To evaluate the prognostic relevance of HistoProfile-derived immune profiles in larger independent datasets, we applied the trained models to H&E WSIs from 11 TCGA cohorts^19^. This analysis revealed cancer type specific associations between disease-specific survival and HistoProfile-estimated biomarkers (**Fig. 6, Supp. Fig. 7**). The strongest positive association was observed for CD8 in HNSC, with a C-index of 0.62 (*P* < 0.001 by permutation test), and HNSC patients with high estimated CD8 density had significantly longer disease-specific survival than patients with low estimated density (HR = 0.49, log-rank *P* < 0.001; **Fig. 6b**). In contrast, estimated CD8 showed strong inverse concordance in KIRC (C-index = 0.32, permutation *P* < 0.001), and patients with high estimated CD8 density had significantly shorter disease-specific survival (HR = 2.72, log-rank *P* < 0.001; **Fig. 6b**).

**Figure 6.**
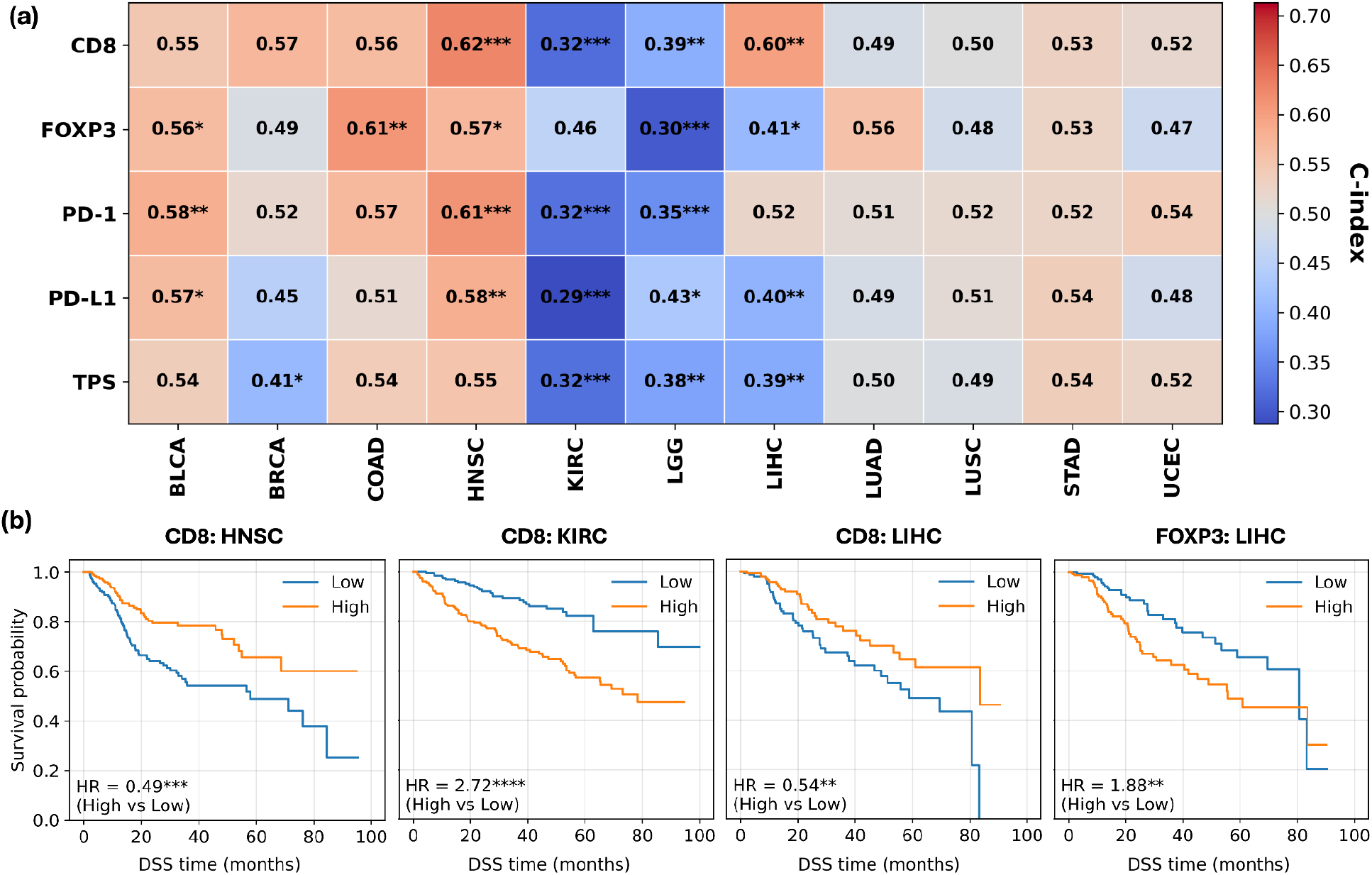
HistoProfile-estimated biomarkers show cancer-specific associations with disease-specific survival in TCGA. **(a)** Disease-specific survival C-indices for HistoProfile-estimated biomarkers across 11 TCGA cancer cohorts. Asterisks indicate permutation-test significance against the null hypothesis of C-index = 0.5: \**P* < 0.05, \*\**P* < 0.01, and \*\*\**P* < 0.001. **(b)** Kaplan-Meier curves illustrating selected cancer-specific survival associations identified in panel (a). Patients were divided into high- and low-biomarker groups using the cohort-specific median estimated value. The HNSC and KIRC curves demonstrate the opposite associations of estimated CD8 density with survival: high estimated CD8 was associated with longer survival in HNSC (HR = 0.49, log-rank *P* < 0.001) but shorter survival in KIRC (HR = 2.72, log-rank *P* < 0.001). In LIHC, high estimated CD8 was associated with longer survival (HR = 0.54, log-rank *P* = 0.005), whereas high estimated FOXP3 was associated with shorter survival (HR = 1.88, log-rank *P* = 0.017).

LIHC further demonstrated opposing prognostic associations between cytotoxic and regulatory immune markers. Estimated CD8 was concordant with longer survival (C-index = 0.60, permutation *P* < 0.01), and patients with high estimated CD8 density had better disease-specific survival (HR = 0.54, log-rank *P* = 0.005). Conversely, estimated FOXP3 was concordant with shorter survival (C-index = 0.41, permutation *P* < 0.05; HR = 1.88, log-rank *P* = 0.017). Altogether, prognostic associations were observed within seven out of the eleven tested cohorts.

## Discussion

We developed HistoProfile, an AI model that estimates key TIME biomarkers (intratumoral CD8, FOXP3, PD-1, and PD-L1 cell densities and PD-L1 TPS) from routine H&E-stained WSIs. Trained using a clinical, pan-cancer cohort with biomarkers quantified via a CLIA-validated mIF assay, the model retains the prognostic stratification provided by the original assay in internal testing and shows survival associations in independent ORION-CRC and TCGA cohorts. By directly estimating established quantitative immune biomarkers, HistoProfile provides an interpretable and scalable link between routine histopathology and clinically validated measurements of the TIME.

An important consideration is whether HistoProfile learns biomarker-specific morphological features or instead relies primarily on general patterns of immune infiltration. Analyses in the held-out ImmunoProfile test set support meaningful biomarker specificity. For each biomarker, HistoProfile estimations were most strongly associated with the corresponding measured biomarker, and although estimations across biomarkers were themselves correlated, these relationships largely recapitulated the correlation structure present in the ground-truth measurements. Moreover, within individual samples, differences between estimated biomarker levels (e.g., estimated CD8 minus FOXP3 density) were positively associated with the corresponding differences in measured biomarker levels. Nonetheless, these analyses are correlative and do not establish the specific morphological features used by the model. Although attention heatmaps provide qualitative insight into regions contributing to estimations, precisely characterizing the histologic features underlying these estimates remains an important direction for future work and a broader challenge in computational pathology.

The TCGA results further highlight the potential of HistoProfile for biomarker exploration in large-scale cohorts where mIF-based quantification is infeasible. The strongest associations with disease-specific survival were observed in KIRC, including a C-index of 0.32 for CD8 estimations. Although CD8-positive lymphocytes are generally considered antitumor effectors, an adverse association between CD8+ T cell infiltration and prognosis is a recognized feature of clear-cell renal cell carcinoma^20–22^. Conversely, the strongest favorable association with survival was observed in HNSC (CD8 C-index of 0.62), consistent with prior analysis of TCGA-HNSC and an independent cohort validated by immunohistochemistry, in which increased cytotoxic T-cell infiltration was associated with improved survival^23^. Although prognostic associations across biomarkers often pointed in the same direction, as also observed in the ImmunoProfile cohort, notable biomarker-specific differences were observed. In LIHC, estimated CD8 and FOXP3 densities showed opposing associations with survival (C-indices of 0.60 and 0.41, respectively), consistent with prior studies describing distinct prognostic roles for intratumoral cytotoxic and regulatory T cell densities in hepatocellular carcinoma^24,25^. Collectively, these findings suggest that HistoProfile-derived biomarker estimates can recover biologically plausible, cancer-specific relationships between the TIME and clinical outcomes.

Our study has several limitations. HistoProfile estimates five key TIME biomarkers and was trained using a large pan-cancer cohort, whereas other mIF assays and corresponding H&E-based estimation approaches have incorporated panels of 20 or more biomarkers, albeit typically with substantially fewer training samples (e.g., tens of slides)^12^. HistoProfile therefore prioritizes sample-level representation and estimation robustness over biomarker breadth. Nonetheless, the included biomarkers capture several major components of the TIME, including cytotoxic and regulatory T-cell populations (CD8 and FOXP3), immune checkpoint states (PD-1 and PD-L1), and tumor-cell PD-L1 expression (TPS). More broadly, the limited availability of large-scale paired H&E/mIF datasets, together with heterogeneity in biomarker staining and quantification protocols across cohorts, creates challenges for external validation. Accordingly, the associations observed in TCGA should be interpreted as correlative evidence of biologically plausible signals rather than definitive characterization of underlying immune biology.

In conclusion, HistoProfile enables scalable estimation of key TIME biomarkers directly from routine H&E histopathology. Across internal and external cohorts, its estimations recapitulate prognostic associations while retaining evidence of biomarker-specific immune information. These findings support HistoProfile as a practical approach for extending standardized immune biomarker profiling to large histopathology cohorts where multiplex assays are unavailable or impractical.

## Methods

### Ethics statement

The IRB of the Dana-Farber Cancer Institute (DFCI) provided ethical approval for this work. All materials were obtained from patients who had signed informed consent forms for the research use of their materials. This research was conducted under DFCI protocols 11–104, 17-000, 20-000, and 22–176.

### Datasets

#### ImmunoProfile cohort

Patients in the ImmunoProfile cohort were enrolled at DFCI between November 2018 and January 2022. Following patient consent, ImmunoProfile testing was ordered at the discretion of the treating clinical team. The assay was available to patients with solid tumors, excluding sarcomas, pediatric malignancies, and neuro-oncology cases. Automated mIF staining was performed using a BOND Rx system (Leica Biosystems, Danvers, MA, USA). The mIF panel comprised five markers: CD8, FOXP3, PD-1, PD-L1, and a tumor-specific marker selected according to cancer type (cytokeratin for epithelial cancers, PAX8 for renal cell carcinomas, and SOX10 for melanomas). DAPI was used as a nuclear counterstain. Stained slides were imaged at 20x magnification using a PhenoImager scanner (Akoya Biosciences, Marlborough, MA, USA). Biomarker quantification was performed over technician-selected ROIs and approved by a pathologist following a predefined SOP. Intratumoral densities of cells positive for each immune biomarker and negative for the tumor marker were computed using a semi-automated image analysis pipeline and approved by a pathologist using a digital sign-out tool. The ImmunoProfile assay and its workflow were validated within the CLIA-certified molecular diagnostics laboratory at the Brigham & Women’s Hospital in Boston (CLIA license #22D2156930). Further details of the assay and quality control can be found in refs^5,16^.

In total, the ImmunoProfile assay was performed on 2,159 samples. For our analysis, we excluded 17 samples without H&E WSIs, and 18 additional samples with insufficient tissue based on manual review, leaving 2,124 samples. Of these, 1,884 were derived from patients represented by a single sample, whereas 116 patients had multiple samples. For patients with multiple samples, a single sample was selected by prioritizing primary biopsies, followed by the earliest available sample, following refs^5,16^. The final cohort comprised 2,000 patients, each represented by one H&E WSI for HistoProfile development. Downstream survival analyses excluded an additional 12 patients with missing follow-up times. The dataset was split into training (1200, 60%), validation (200, 10%), and test (600, 30%) sets while stratifying for cancer types and binarized cancer stage (0, stages I–II; 1, stages III–IV).

#### ORION-CRC cohort

The ORION-CRC cohort consists of 40 colorectal cancer resection specimens from patients with stage II-IV disease, collected from the Brigham and Women’s Hospital pathology archives^18^. Each specimen was imaged as a whole slide using the ORION platform, which acquired mIF measurements followed by H&E staining and imaging of the same tissue section. The ORION-CRC cohort was reserved for held-out validation in the current study. Cell densities for CD8, FOXP3, PD-1, and PD-L1 and PD-L1 TPS were computed over the inner tumor (“R1”) region using the pre-processed cell table for each sample to maximize alignment with ImmunoProfile biomarker measurement definitions.

#### TCGA cohort

Diagnostic H&E WSIs of primary solid tumors were analyzed. Patients missing either disease-specific survival status or disease-specific survival time were excluded. When multiple eligible slides remained for the same patient, one slide was selected at random, yielding one diagnostic slide per patient. To ensure adequate cohort size and sufficient numbers of survival events, downstream analyses were restricted to cancer cohorts containing at least 300 eligible patients and at least 50 disease-specific deaths.

#### H&E embedding generation

H&E WSIs for each dataset were processed using TRIDENT (v0.2.3)^26^ and PathFMTools^27^. Tissue-containing regions were identified using the HEST tissue segmentation model, and patches of 224 x 224 pixels were extracted from the segmented tissue regions at 20x magnification without overlap. Embeddings were generated for the extracted patches using CONCH v1^28^ with the encoder weights held fixed during downstream model training and testing, producing a 512-dimensional feature representation. The resulting patch-level embeddings were retained for downstream attention-based multiple-instance learning.

### HistoProfile model and training

Each WSI was represented as a bag of patch embeddings 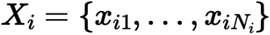,where **x**_*ij*_=ℝ^*E*^ is the feature vector for patch *j* in slide *i*. The CONCH v1^28^ foundation model was used for embedding generation, with embedding size of *E* = 512. The learning target *y*_*i*_ was set to a percentile-normalized immune biomarker measurement in [0,1]. Percentiles for normalization were computed using the training set.

We trained a weakly supervised attention-based multiple instance learning (ABMIL^17^) model to aggregate patch embeddings into a slide-level representation. For each patch Embedding **x**_*ij*_, a gated attention score was computed as *w*_*ij*_=**w**^⊤^(tanh(**W**_*u*_**x**_*ij*_) ⊙*σ*(**W**_*v*_**x**_*ij*_)), with **W**_*u*_,**W**_*v*_∈ℝ^H⨯*E*^, hidden dimension *H*, sigmoid *σ*(⋅), and elementwise product ⊙. Attention weights were normalized within each slide, 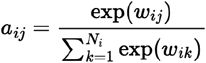, and used to form a slide embedding 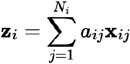. A linear prediction 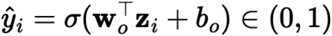 head produced a scalar prediction. Dropout was applied to the attention projections during training.

Models were trained with mean absolute error (MAE) using Adam^29^ optimization (batch size *B*=1 slide per update due to variable bag sizes). Learning rate *η* was set to 1e-3. Attention hidden dimension and dropout rate were tuned via grid search over values of (64, 128, 256) and (0.1, 0.3, 0.5) respectively, and model selection used the checkpoint with minimum validation loss. A separate model was trained for each biomarker. Models were trained for a maximum of 500 epochs, with early stopping based on validation loss (patience of 15 epochs). The chosen set of hyperparameters (number of epochs, dropout rate, hidden dimension) for each biomarker were as follows: (27, 0.1, 128) for CD8, (14, 0.1, 256) for FOXP3, (7, 0.1, 128) for PD-L1, (28, 0.1, 64) for PD-1, and (16, 0.1, 256) for TPS.

### Downstream survival analysis

For pan-cancer analysis in the ImmunoProfile cohort, overall survival associations were evaluated using Cox proportional hazards regression stratified by dichotomized cancer stage (0, stages I–II; 1, stages III–IV) and cancer type (non-small cell lung cancer, breast cancer, colorectal cancer, or other), following previous analysis in this dataset. Subgroup analyses by cancer type were stratified by dichotomized cancer stage only. For ORION-CRC and TCGA, C-indices were calculated directly between HistoProfile-estimated biomarker values and observed survival outcomes.

### Statistical analysis

Statistical significance for the Pearson correlation between measured and estimated biomarker values was evaluated using a two-sided t-test with the null hypothesis of zero correlation. Confidence intervals for correlation coefficients and C-indices were estimated using nonparametric bootstrapping with 5,000 resamples. Statistical significance for C-indices was assessed using a two-sided permutation test of the null hypothesis that the C-index was 0.5. For the Kaplan-Meier curves, the log-rank test compared the observed and expected numbers of events between groups across event times. Statistical significance was determined using the asymptotic chi-squared distribution of the log-rank statistic with (k-1) degrees of freedom, where k denotes the number of groups compared.

## Data Availability

The TCGA data analyzed in this study are publicly available through the National Cancer Institute Genomic Data Commons (https://portal.gdc.cancer.gov/). The ORION-CRC dataset is publicly available through Zenodo at https://zenodo.org/records/7637988. The ImmunoProfile dataset is not publicly available because it contains data derived from human participants and is subject to institutional and patient privacy restrictions.

