## Supplementary Information for "Pan-cancer estimation of prognostic immune biomarkers from histopathology using artificial intelligence"

| | | Stage | | | Tissue Sample Type | | | Split | | |
|---|---|---|---|---|---|---|---|---|---|---|
| **Cancer Type** | **Count (%)** | **Low** | **High** | **Unknown** | **Primary** | **Metastatic** | **Unknown** | **Train** | **Val.** | **Test** |
| **Non-Small Cell Lung Cancer** | 504 (25.2) | 302 (59.9) | 199 (39.5) | 3 (0.6) | 412 (81.7) | 87 (17.3) | 5 (1.0) | 302 (59.9) | 50 (9.9) | 152 (30.2) |
| **Breast Cancer** | 247 (12.4) | 143 (57.9) | 104 (42.1) | 0 (0.0) | 190 (76.9) | 54 (21.9) | 3 (1.2) | 148 (59.9) | 24 (9.7) | 75 (30.4) |
| **Colorectal Cancer** | 230 (11.5) | 47 (20.4) | 183 (79.6) | 0 (0.0) | 128 (55.7) | 101 (43.9) | 1 (0.4) | 138 (60.0) | 23 (10.0) | 69 (30.0) |
| **Bladder Cancer** | 160 (8.0) | 99 (61.9) | 61 (38.1) | 0 (0.0) | 139 (86.9) | 21 (13.1) | 0 (0.0) | 96 (60.0) | 16 (10.0) | 48 (30.0) |
| **Esophagogastric Cancer** | 122 (6.1) | 35 (28.7) | 84 (68.9) | 3 (2.5) | 91 (74.6) | 30 (24.6) | 1 (0.8) | 73 (59.8) | 12 (9.8) | 37 (30.3) |
| **Melanoma** | 108 (5.4) | 23 (21.3) | 84 (77.8) | 1 (0.9) | 42 (38.9) | 65 (60.2) | 1 (0.9) | 65 (60.2) | 11 (10.2) | 32 (29.6) |
| **Pancreatic Cancer** | 104 (5.2) | 31 (29.8) | 73 (70.2) | 0 (0.0) | 49 (47.1) | 55 (52.9) | 0 (0.0) | 62 (59.6) | 10 (9.6) | 32 (30.8) |
| **Head and Neck Cancer** | 94 (4.7) | 38 (40.4) | 55 (58.5) | 1 (1.1) | 76 (80.9) | 18 (19.1) | 0 (0.0) | 56 (59.6) | 10 (10.6) | 28 (29.8) |
| **Endometrial Cancer** | 92 (4.6) | 54 (58.7) | 38 (41.3) | 0 (0.0) | 75 (81.5) | 17 (18.5) | 0 (0.0) | 55 (59.8) | 9 (9.8) | 28 (30.4) |
| **Ovarian Cancer** | 91 (4.6) | 18 (19.8) | 72 (79.1) | 1 (1.1) | 42 (46.2) | 49 (53.8) | 0 (0.0) | 55 (60.4) | 9 (9.9) | 27 (29.7) |
| **Renal Cell Carcinoma** | 83 (4.2) | 23 (27.7) | 60 (72.3) | 0 (0.0) | 43 (51.8) | 40 (48.2) | 0 (0.0) | 50 (60.2) | 8 (9.6) | 25 (30.1) |
| **Hepatobiliary Cancer** | 34 (1.7) | 12 (35.3) | 21 (61.8) | 1 (2.9) | 27 (79.4) | 7 (20.6) | 0 (0.0) | 21 (61.8) | 4 (11.8) | 9 (26.5) |
| **Prostate Cancer** | 34 (1.7) | 7 (20.6) | 27 (79.4) | 0 (0.0) | 18 (52.9) | 16 (47.1) | 0 (0.0) | 21 (61.8) | 4 (11.8) | 9 (26.5) |
| **Small Cell Lung Cancer** | 20 (1.0) | 5 (25.0) | 15 (75.0) | 0 (0.0) | 12 (60.0) | 7 (35.0) | 1 (5.0) | 12 (60.0) | 2 (10.0) | 6 (30.0) |
| **Other** | 77 (3.8) | 4 (5.2) | 35 (45.5) | 38 (49.4) | 37 (48.1) | 32 (41.6) | 8 (10.4) | 46 (59.7) | 8 (10.4) | 23 (29.9) |
| **Total** | 2000 (100.0) | 841 (42.0) | 1111 (55.6) | 48 (2.4) | 1381 (69.0) | 599 (30.0) | 20 (1.0) | 1200 (60.0) | 200 (10.0) | 600 (30.0) |

**Supplementary Table 1.** ImmunoProfile cohort summary features by cancer type, stage, tissue sample type, and training, validation, and test split.

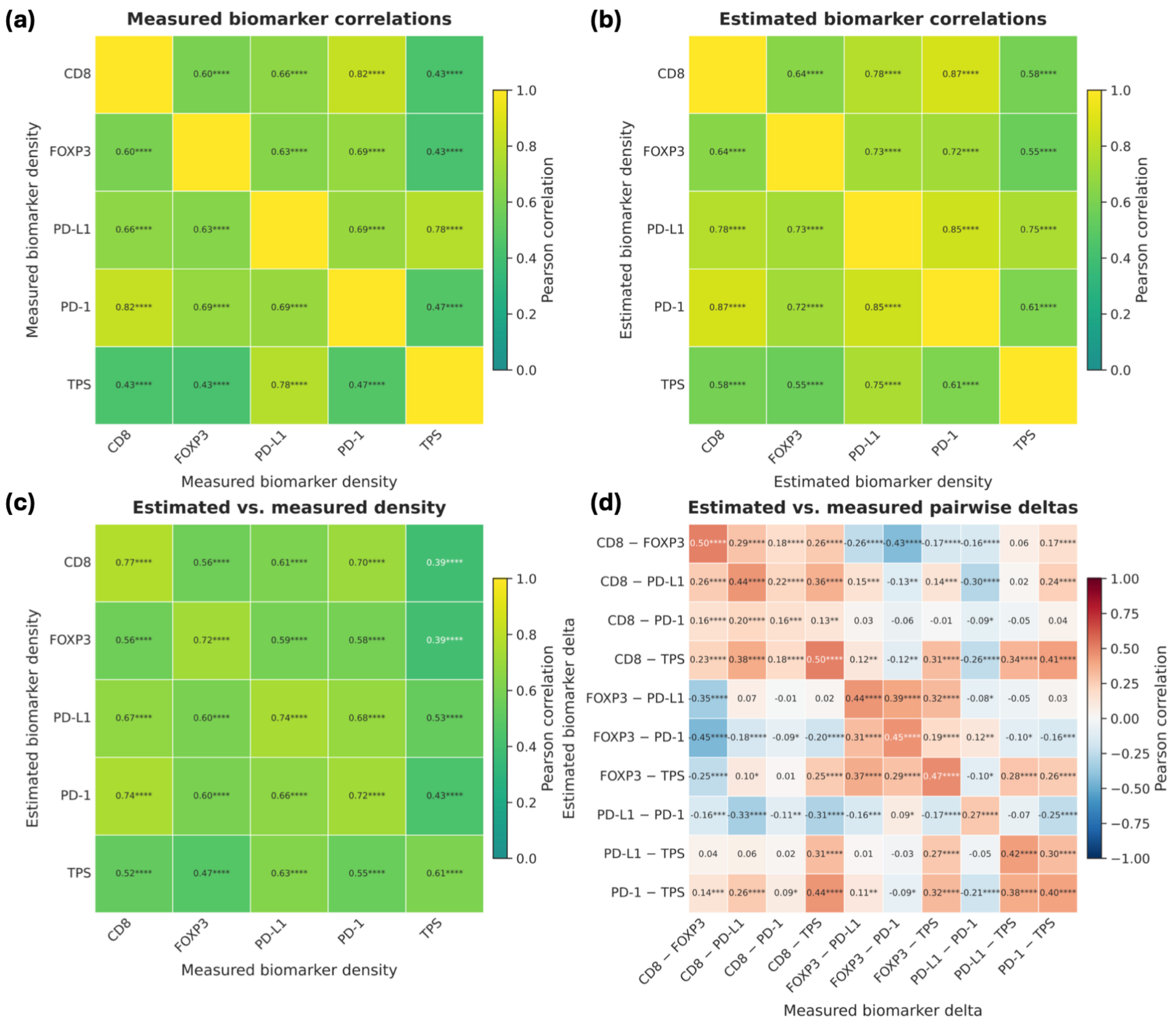


**Supplementary Figure 1. HistoProfile estimates immune biomarkers with high correlation to the measured values and distinguishes between biomarkers. (a)** Pearson correlation between measured biomarker pairs. **(b)** Pearson correlation between estimated biomarker pairs. **(c)** Pearson correlation between estimated and measured biomarker values. **(d)** Pearson correlation between estimated and measured difference between pairs of immune biomarkers. (* $P < 0.05$, ** $P < 0.01$, *** $P < 0.001$, and **** $P < 0.0001$).

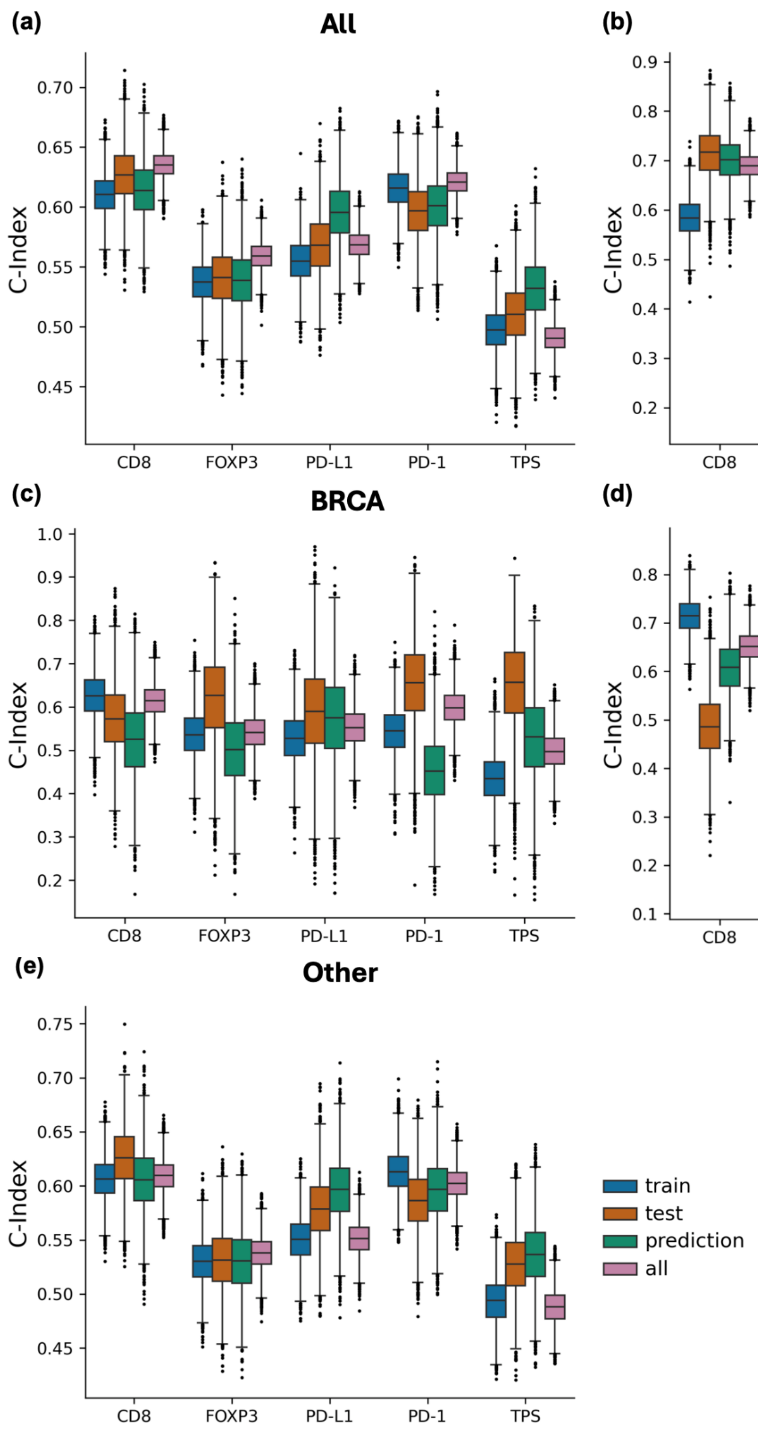


**Supplementary Figure 2. Comparison of biomarker C-indices in ImmunoProfile data splits.** Bootstrapped C-indices (5000 bootstrap resamples) of measured biomarkers in training, testing, and combined data splits and HistoProfile-estimated values for (a) the pan-cancer ImmunoProfile cohort ("All"), and separately for (b) non-small cell lung cancer (NSCLC), (c) breast cancer (BRCA), (d) colorectal cancer (CRC), and (e) other cancer types.

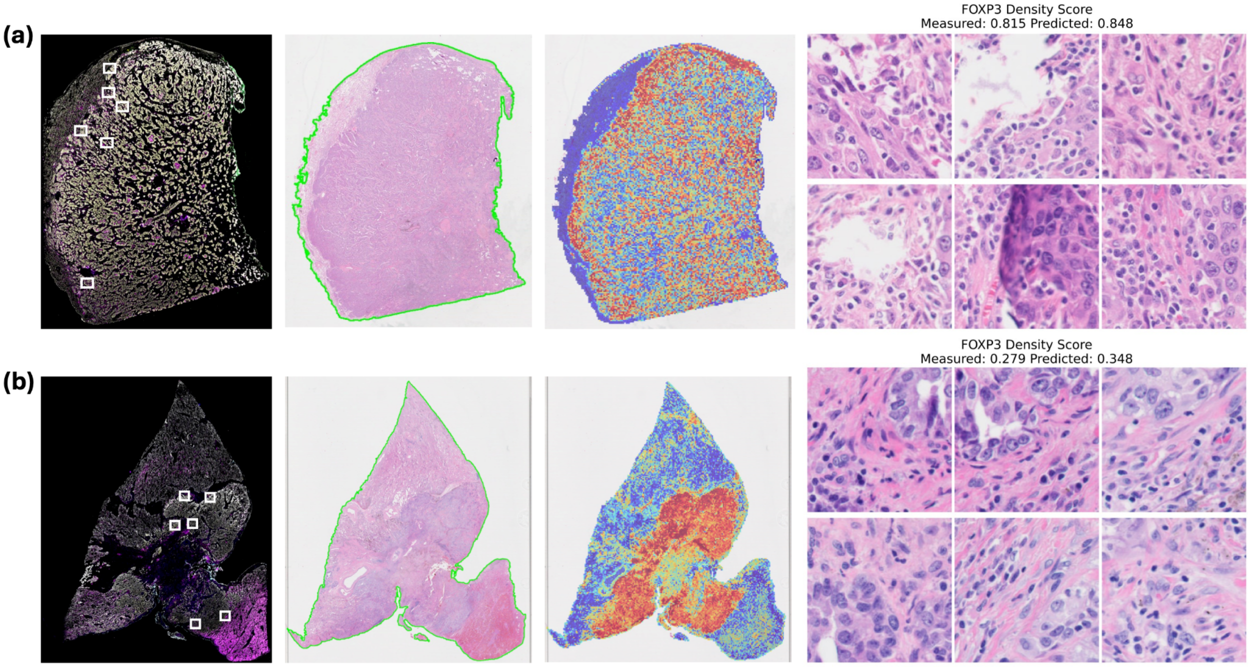


**Supplementary Figure 3. Visualization of HistoProfile attention patterns for representative samples with high and low FOXP3 density. (a, b)** Representative samples with high (a) and low (b) normalized FOXP3 density scores. From left to right, each row shows the multiplex immunofluorescence (mIF) whole-slide image with expert-selected regions of interest indicated by white boxes, the corresponding H&E whole-slide image with the analyzed tissue region outlined, the HistoProfile attention map, and the most highly attended H&E patches. Warmer colors in the attention map indicate higher model attention. Measured and HistoProfile-estimated FOXP3 density scores are reported above the patch panels. Both examples are from NSCLC primary tumor specimens.

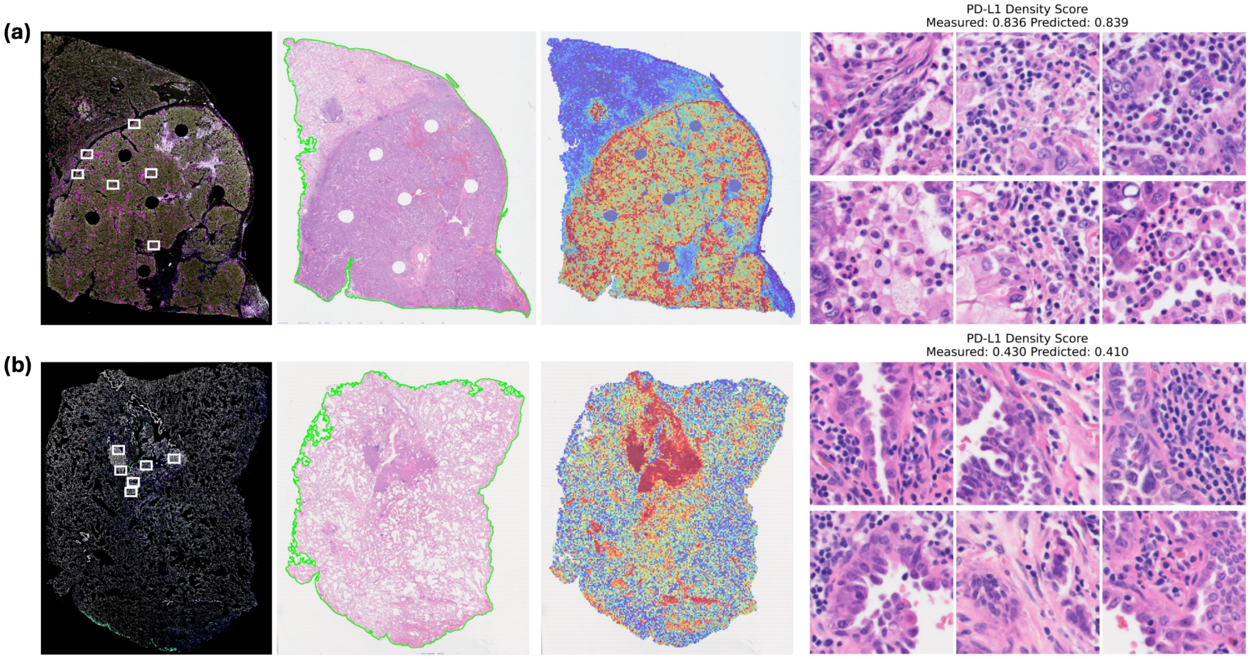


**Supplementary Figure 4. Visualization of HistoProfile attention patterns for representative samples with high and low PD-L1 density.** (a, b) Representative samples with high (a) and low (b) normalized PD-L1 density scores. From left to right, each row shows the multiplex immunofluorescence (mIF) whole-slide image with expert-selected regions of interest indicated by white boxes, the corresponding H&E whole-slide image with the analyzed tissue region outlined, the HistoProfile attention map, and the most highly attended H&E patches. Warmer colors in the attention map indicate higher model attention. Measured and HistoProfile-estimated PD-L1 density scores are reported above the patch panels. Both examples are from NSCLC primary tumor specimens.

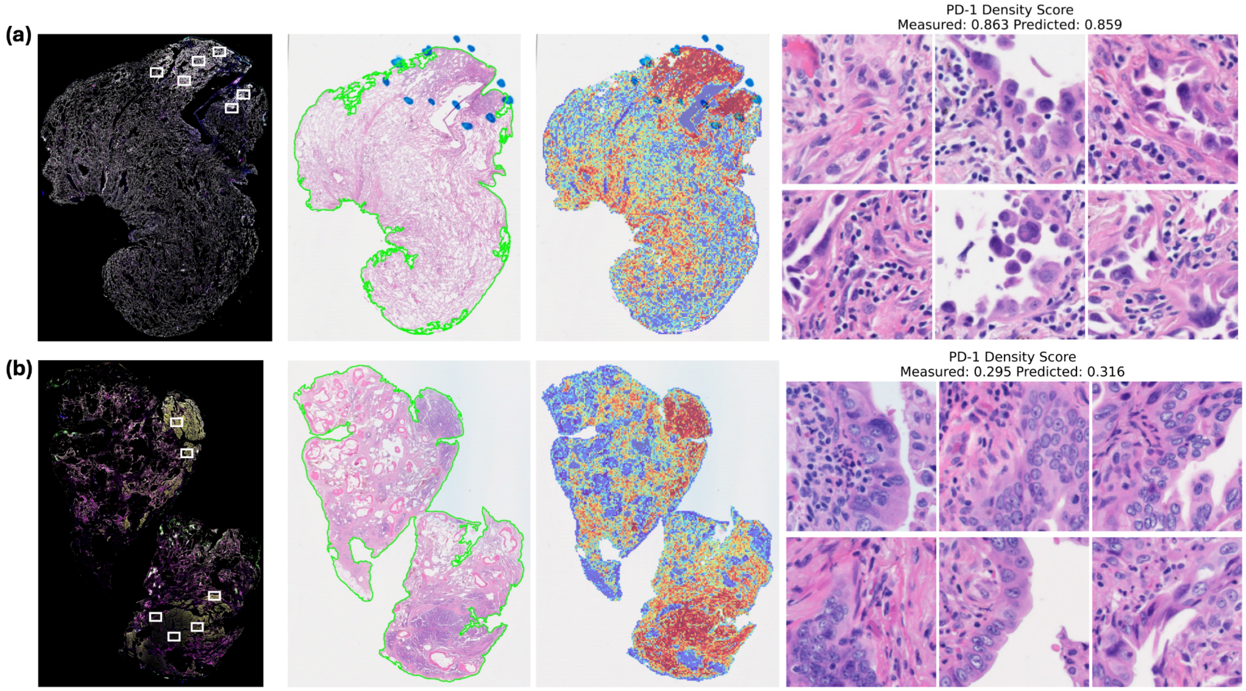


**Supplementary Figure 5. Visualization of HistoProfile attention patterns for representative samples with high and low PD-1 density.** (a, b) Representative samples with high (a) and low (b) normalized PD-1 density scores. From left to right, each row shows the multiplex immunofluorescence (mIF) whole-slide image with expert-selected regions of interest indicated by white boxes, the corresponding H&E whole-slide image with the analyzed tissue region outlined, the HistoProfile attention map, and the most highly attended H&E patches. Warmer colors in the attention map indicate higher model attention. Measured and HistoProfile-estimated PD-1 density scores are reported above the patch panels. Both examples are from NSCLC primary tumor specimens.

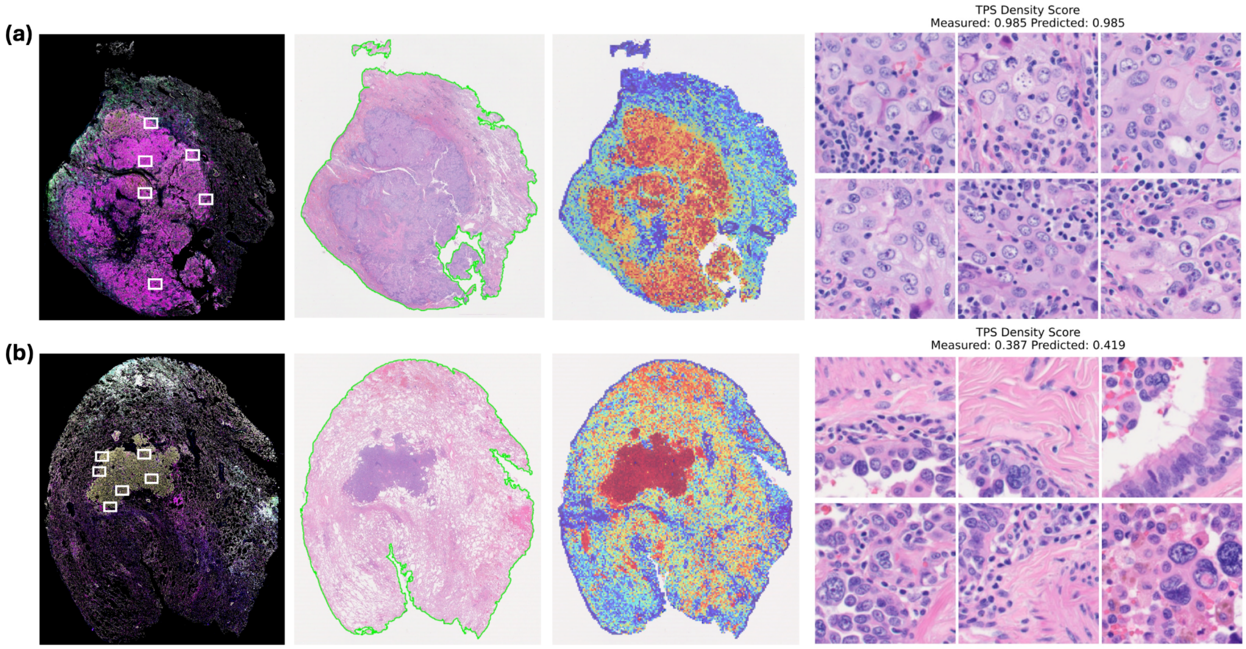


**Supplementary Figure 6. Visualization of HistoProfile attention patterns for representative samples with high and low TPS.** (a, b) Representative samples with high (a) and low (b) normalized TPS scores. From left to right, each row shows the multiplex immunofluorescence (mIF) whole-slide image with expert-selected regions of interest indicated by white boxes, the corresponding H&E whole-slide image with the analyzed tissue region outlined, the HistoProfile attention map, and the most highly attended H&E patches. Warmer colors in the attention map indicate higher model attention. Measured and HistoProfile-estimated TPS scores are reported above the patch panels. Both examples are from NSCLC primary tumor specimens.

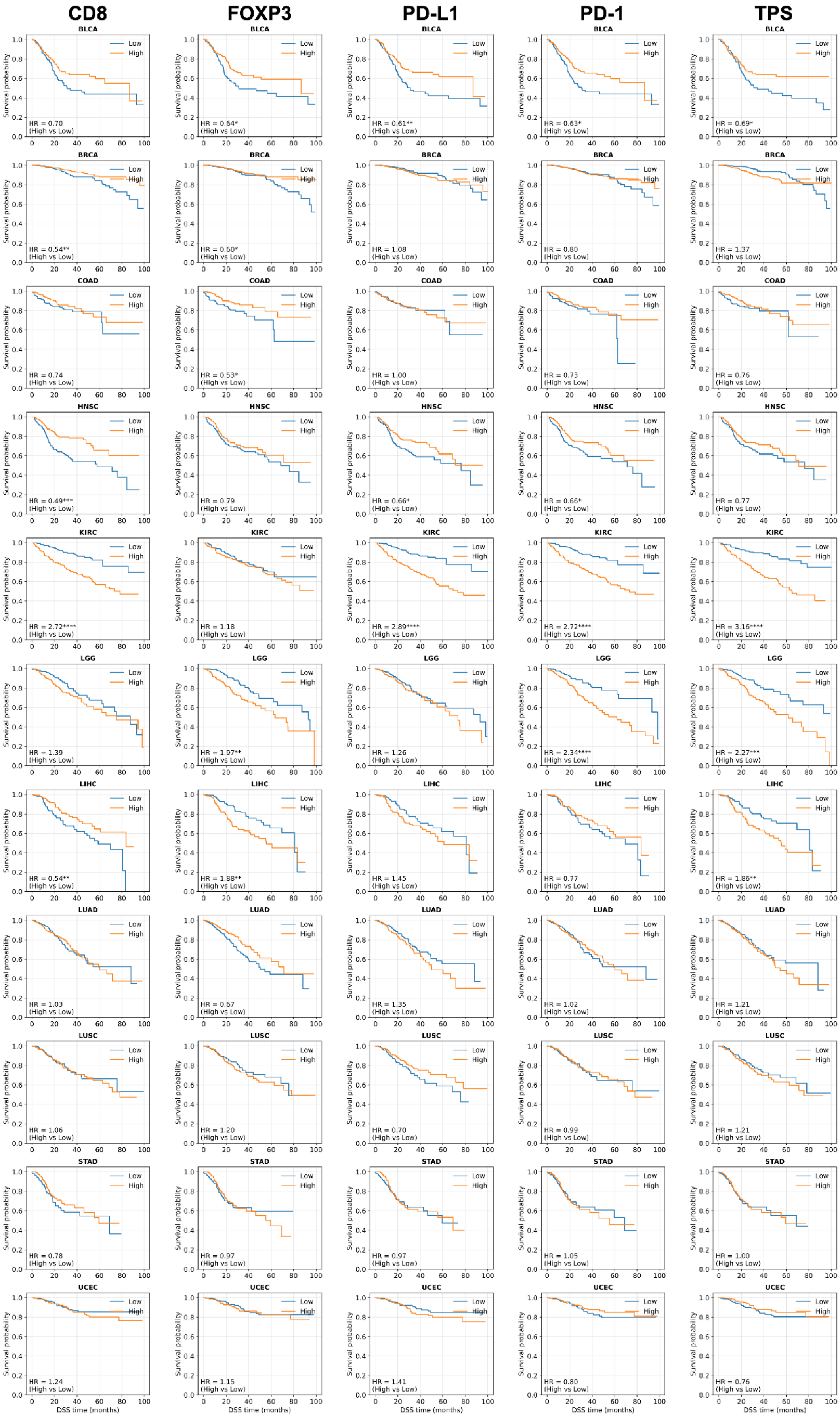
CD8
FOXP3
PD-L1
PD-1
TPS
BLCA
HR = 0.70 (High vs Low)
HR = 0.64* (High vs Low)
HR = 0.61** (High vs Low)
HR = 0.63* (High vs Low)
HR = 0.69* (High vs Low)
BRCA
HR = 0.54** (High vs Low)
HR = 0.60* (High vs Low)
HR = 1.08 (High vs Low)
HR = 0.80 (High vs Low)
HR = 1.37 (High vs Low)
COAD
HR = 0.74 (High vs Low)
HR = 0.53* (High vs Low)
HR = 1.00 (High vs Low)
HR = 0.73 (High vs Low)
HR = 0.76 (High vs Low)
HNSC
HR = 0.49*** (High vs Low)
HR = 0.79 (High vs Low)
HR = 0.66* (High vs Low)
HR = 0.66* (High vs Low)
HR = 0.77 (High vs Low)
KIRC
HR = 2.72**** (High vs Low)
HR = 1.18 (High vs Low)
HR = 2.89**** (High vs Low)
HR = 2.72**** (High vs Low)
HR = 3.16**** (High vs Low)
LGG
HR = 1.39 (High vs Low)
HR = 1.97** (High vs Low)
HR = 1.26 (High vs Low)
HR = 2.34**** (High vs Low)
HR = 2.27*** (High vs Low)
LIHC
HR = 0.54** (High vs Low)
HR = 1.88** (High vs Low)
HR = 1.45 (High vs Low)
HR = 0.77 (High vs Low)
HR = 1.86** (High vs Low)
LUAD
HR = 1.03 (High vs Low)
HR = 0.67 (High vs Low)
HR = 1.35 (High vs Low)
HR = 1.02 (High vs Low)
HR = 1.21 (High vs Low)
LUSC
HR = 1.06 (High vs Low)
HR = 1.20 (High vs Low)
HR = 0.70 (High vs Low)
HR = 0.99 (High vs Low)
HR = 1.21 (High vs Low)
STAD
HR = 0.78 (High vs Low)
HR = 0.97 (High vs Low)
HR = 0.97 (High vs Low)
HR = 1.05 (High vs Low)
HR = 1.00 (High vs Low)
UCEC
HR = 1.24 (High vs Low)
HR = 1.15 (High vs Low)
HR = 1.41 (High vs Low)
HR = 0.80 (High vs Low)
HR = 0.76 (High vs Low)
Low
High
Survival probability
DSS time (months)

**Supplementary Figure 7. HistoProfile estimated immune biomarkers show prognostic characteristics with respect to disease-specific survival in TCGA.** Kaplan-Meier survival curves for 11 cancer types based on HistoProfile-estimated CD8, FOXP3, PD-L1, PD-1, and TPS values. High and low groups were defined using the respective cohort medians (* $P < 0.05$, ** $P < 0.01$, *** $P < 0.001$, and **** $P < 0.0001$).
